# Differential effectiveness of bee and beetle pollinators creates flower colour polymorphism

**DOI:** 10.64898/2026.01.12.698860

**Authors:** Jonathan Heinze, Casper J. van der Kooi, Johannes Spaethe

## Abstract

Floral diversity is the hallmark of Angiosperms and is often shaped by pollinator-mediated selection, which drives floral divergence as plants evolve traits that attract specific pollinator groups. Mediterranean red, bowl-shaped flowers are specialized on pollination by glaphyrid beetles in areas with beetles and bees but evolved different colours and specialize on bees as pollinators when beetles are absent. What explains this geographic variation in specialization and concomitant flower colour differences?

We assessed pollination effectiveness of honeybees and glaphyrids for the colour-polymorphic *Anemone pavonina* in its natural habitat. We recorded the duration and number of flower visits and quantified single-visit seed set. Per-visit effectiveness (relative seed set) was threefold higher for *Pygopleurus* (36.3%) than for honeybees (9.5%). Flower visitation lasted substantially longer for beetles (56.3 sec. on average) than for honeybees (2.3 sec.). For beetles, pollinator importance (single-visit seed set x visitation frequency) was highest at low elevations but declined with increasing elevation, whereas for bees pollinator importance remained low yet constant across the elevational gradient.

The high pollinator effectiveness of beetles indicates that floral specialization for this pollinator group is advantageous, even to the point that excluding low-quality bee pollinators – by producing inconspicuous flower colours – is beneficial. Non-red flowers may only be beneficial at higher elevations, where bee pollinator importance exceeds that of beetles.

## Introduction

The floral diversity of angiosperms epitomizes how natural selection generates phenotypic variation. Pollinator-mediated selection is a key driver of this process [1]. Geographic variation in pollinator abundance can cause intraspecific floral trait variation that aligns with the local pollinator community [2, 3].

The European flora hosts an enormous diversity in shapes, sizes and colours of flowers, yet red-flowered species are exceptionally rare throughout continental Europe [4]. Across many continents, red flowers are associated with pollination by birds, which see red colours whereas many insects do not [5]; however, bird pollination is virtually absent in Europe [6]. In the Eastern Mediterranean, there is a remarkable case of convergent evolution of red, bowl-shaped flowers, which have repeatedly evolved in at least six genera: *Anemone, Adonis* and *Ranunculus* (Ranunculaceae), *Papaver* and *Glaucium* (Papaveraceae), and *Tulipa* (Liliaceae). This floral syndrome is referred to as the so-called ‘poppy guild’ [7, 8]. Glaphyrid beetles (*Pygopleuruss* spp.) can see red colours [9], use these flowers as mating sites and food sources, and are considered the primary pollinators of this guild [7, 8, 10].

Several species in the poppy guild (e.g. *Papaver, Ranunculus* and *Anemone* spp.) exhibit intra-specific flower colour polymorphism. Dark red morphs are beetle pollinated and ultraviolet reflecting or purple morphs are pollinated by bees and occasionally flies [8, 11, 12]. Intriguingly, these bees also occur in regions where red-flowered morphs occur. In other words, in areas where bees and red-sensitive beetles co-occur, poppy guild flowers are specialized for beetle pollination and have colours that are inconspicuous to bees, whereas in areas where these beetles are absent flowers are pollinated by bees and have different colours [13].

What explains the apparent specialization of poppy guild flowers to beetles whenever they are present, yet enables pollination by bees as secondary pollinators when necessary? The “Most Effective Pollinator Principle” [14] predicts that the characteristics of flowers at a certain location are “molded” by the most effective pollinator [see also 15, 16]. Consequently, when beetles are better pollinators than bees, it will be advantageous for the plant to exclude bees (e.g. via inconspicuous floral colours).

*Anemone pavonina* (Ranunculaceae) exhibits red and purple flowering morphs along an elevational gradient on the slopes of Mount Olympus in Greece. Red flowers occur at lower elevations and are associated with glaphyrid beetle pollination, whereas purple flowers grow at higher elevations and are associated with bee pollination [12]. It has been suggested that the upper limit of red flowers at ca. 1200 m a.s.l. is also the distribution limit of *Pygopleurus* beetles [12, 13]. Bees that are common at both low and high elevations, strongly prefer purple over red flowers [13], presumably because bees have very minimal sensitivity to long-wavelength ranges [17].

Here, we quantify pollination performance of bees and beetles on *A. pavonina* in its native range. We measured pollinator effectiveness (quantified as relative seed set per single-visit) of honeybees (*Apis mellifera*) and the most common glaphyrid beetle (*Pygopleurus chrysonotus*) on red *A. pavonina* flowers in the field. Pollinator importance was calculated as pollinator effectiveness x visitation frequency [18-20]. We hypothesized that beetles are more effective pollinators than bees and therefore enhance the reproductive success of the plants. Consequently, in areas with sufficiently high *Pygopleurus* abundance, beetle pollinator importance exceeds that of bees, making it beneficial for flowers to specialize on beetle pollinators (i.e. evolve red flower colours).

## Material and Methods

### Study site, plant species and pollinators

The experiments were performed during flowering season between March and May in Thessaly, northern Greece. At the southern slopes of Mt. Olympus, *Anemone pavonina* can be found in large monomorphic red populations at lower elevations (500-1000 m a.s.l), in purple populations at higher elevations (1200-1500 m a.s.l), and in a narrow transition zone of colour polymorphic populations in between (1000-1200 m a.s.l). *Anemone pavonina* only offers pollen as nutritional reward. Glaphyrid beetles (*Pygopleurus chrysonotus*), used for the pollination experiments in netted tents, were collected in the evening from flowers where they typically shelter overnight. The beetles were kept in a net cage with *Anemone* flowers as food source for at least 12 hours prior to the experiments. Experiments with bees were performed with honeybee (*Apis mellifera*) foragers in the field at sites where they were common. We observed honeybees collecting pollen from *A. pavonina* flowers of both colours, which allowed us to directly compare beetle and bee pollination performance on the same colour morph. We also observed various wild bee species foraging on anemones [13]; however, because of their low abundance, we focused on honeybees as a proxy for bee pollination in general [21].

### Pollination experiments

To assess pollinator effectiveness (relative seed set per visit), we collected plants of the red morph from monomorphic *A. pavonina* populations and potted them individually in commercially available brown plastic pots (ø 14 cm). We bagged all buds with small mesh bags (12x15cm, mesh width ca. 0.55 x 0.35mm) and labelled the flowers individually. For testing honeybees, we placed individually potted plants (with only one recipient flower per individual to avoid geitonogamous pollination) from a different source population within a polymorphic *A. pavonina* population and waited for honeybees to visit a flower (Figure 1A). We counted only visits when honeybees had visited several other anemone flowers before.

**Figure 1:**
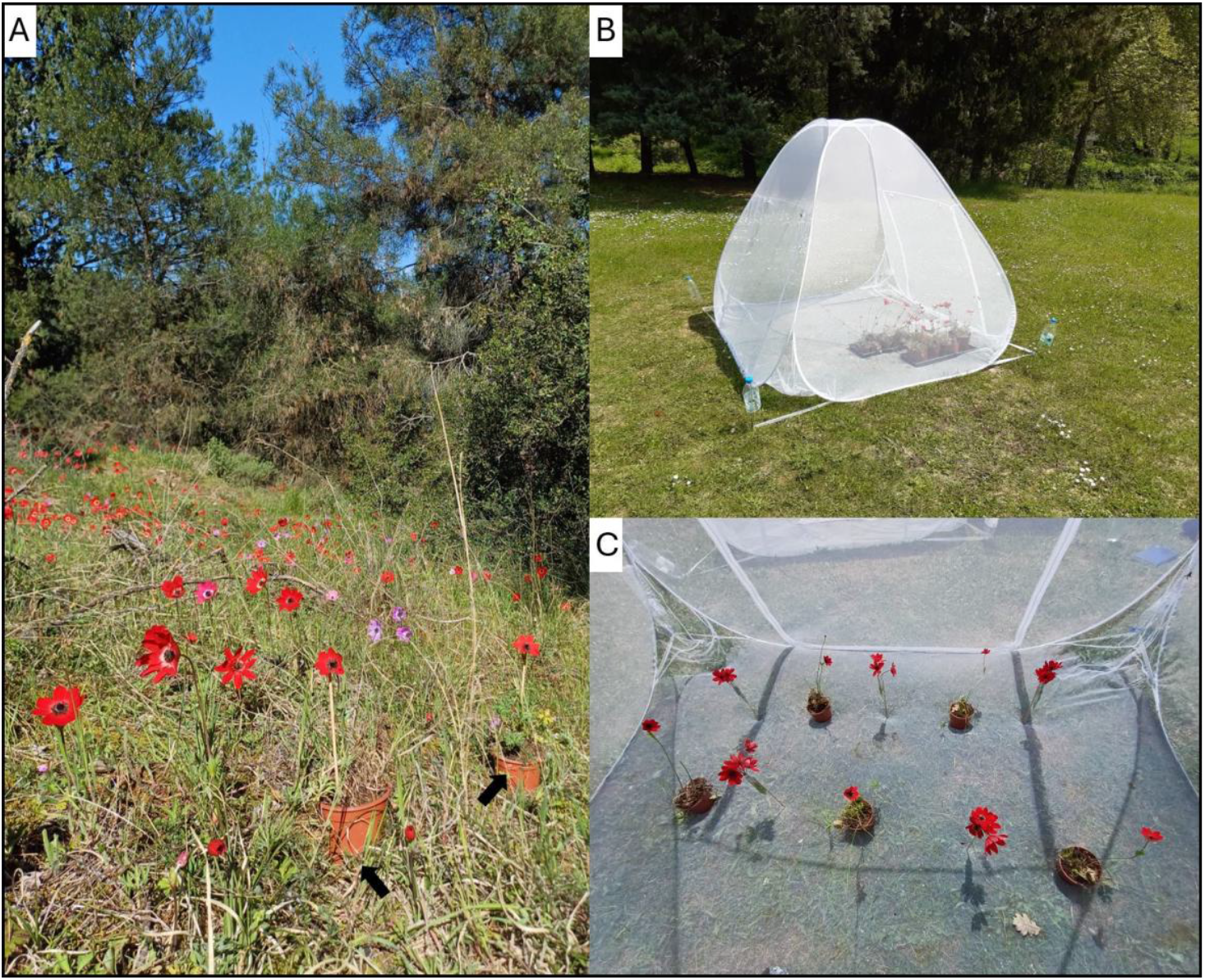
(A) Natural population of *A. pavonina* with interspersed, potted plants from the pollination experiment (black arrows). (B) Mesh tent setup for the beetle pollination experiment and (C) arrangement of potted receiver-flowers and donor-flowers in small vials within a 2 x 5 checkerboard array.

The beetle experiment took place in a white netted tent (Figure 1B; 2 m x 1.8 m, height 1.6 m, mesh 300/ inch^2^), close to the anemone population where the honeybees were tested. During the experiment, we presented five potted plants (again with one recipient flower each) that had opened within the last 24-48 hours and five vials with male-phase, pollen donor flowers. The flowers were arranged in a 2 x 5 checkerboard array (Figure 1C). We then placed 5-10 beetles on the donor flowers, where they immediately started feeding on pollen. All beetles visited at least two to three donor flowers for a minimum of five minutes in total and thus had had ample contact with stamens and pollen, before they approached the receiver flower.

We recorded the time a beetle or honeybee spent on the flower from touching the stigmas until it left. We bagged the flower as soon as the insect had left to prevent further pollination. In natural populations flowers may receive more than one pollinator visit, but the duration of a single visit varies considerably (0.5 sec to 45 min). We thus accounted for the highly variable visit duration by allowing one or two visits per flower and took the total duration of both and calculated the average time per visit. As a negative control to quantify autonomous self-pollination, we bagged flowers to prevent insect pollination. To control for maximum seed set, we hand pollinated another group of flowers with a small paint brush (round brush, synthetic bristles, size 10) with pollen of three other flowers. The receptivity of a flower during the female phase decreases with age [13], so we only used flowers up to two days after the flowers had opened. Pollinated flowers were kept separately in an outside area and were watered every day. Infructescences (seed heads) were collected after about two weeks when they were dry and starting to fall apart. Relative seed set was calculated as the total number of carpels (there is only one ovule per carpel in *Anemone*) relative to developed seeds (number of seeds/number of carpels). Pollinator importance [18, 19] was calculated as the product of pollinator effectiveness (seed set per visit) and visitation frequency (visits per hour) to quantify spatiotemporal patterns of pollinator performance [20]. Pollinator visitation frequency was estimated based on insect-trapping data from our study region [13], which we extended with data obtained in 2025.

### Statistics

Statistical analyses were performed in R (Version 4.2.1 2022-06-23) and the additional packages “rstatix” [22] and “dplyr” [23]. To compare relative seed set between the four different groups (pollinator exclusion, beetle pollination, honeybee pollination and hand pollination) and between duration per flower visit of *Pygopleurus* and honeybees, we used a Kruskal-Wallis test due to non-normally distributed residuals and variance inhomogeneity in our data, followed by a Dunn post-hoc test with “holm” correction for multiple testing. To model pollinator importance along the elevational gradient, we used a generalized linear model (glm) with pollinator importance as response and elevation as predictor variable.

## Results

We tested pollination effectiveness of 31 honeybees on 19 flowers and 18 beetles on 12 flowers. Beetles had a higher per-visit effectiveness than honeybees (Figure 2). The Kruskal-Wallis test showed significant differences between the four pollination treatments (p<0.001; df =3; χ^2^=46.607). The average relative seed set of flowers pollinated by beetles was 36.3% ± 6 (mean ± SE), which is significantly higher (p=0.038) than the 9.5% ± 2 relative seed set of plants pollinated by honeybees. For both pollinator groups, relative seed set was higher (honeybees: p=0.026; beetles: p<0.001) than for the pollinator exclusion control (0.5% ± 0.4) but less than for the hand pollination treatment (79.3% ± 3.2; honeybees: p<0.001; beetles: p=0.046). Beetles were found to spend (much) more time per-visit on flowers than bees (p<0.001; df=1; χ^2^=14.66; beetles: mean 56.3 seconds; honeybees: mean 2.3 seconds). Pollinator importance decreased with elevation for beetles (p = 0.018), but not for honeybees (p = 0.124). At low elevations (around 500m a.s.l.) pollinator abundance was approximately ten times higher for beetles than for bees but decreased to a similar level, and below, at high elevations.

**Figure 2:**
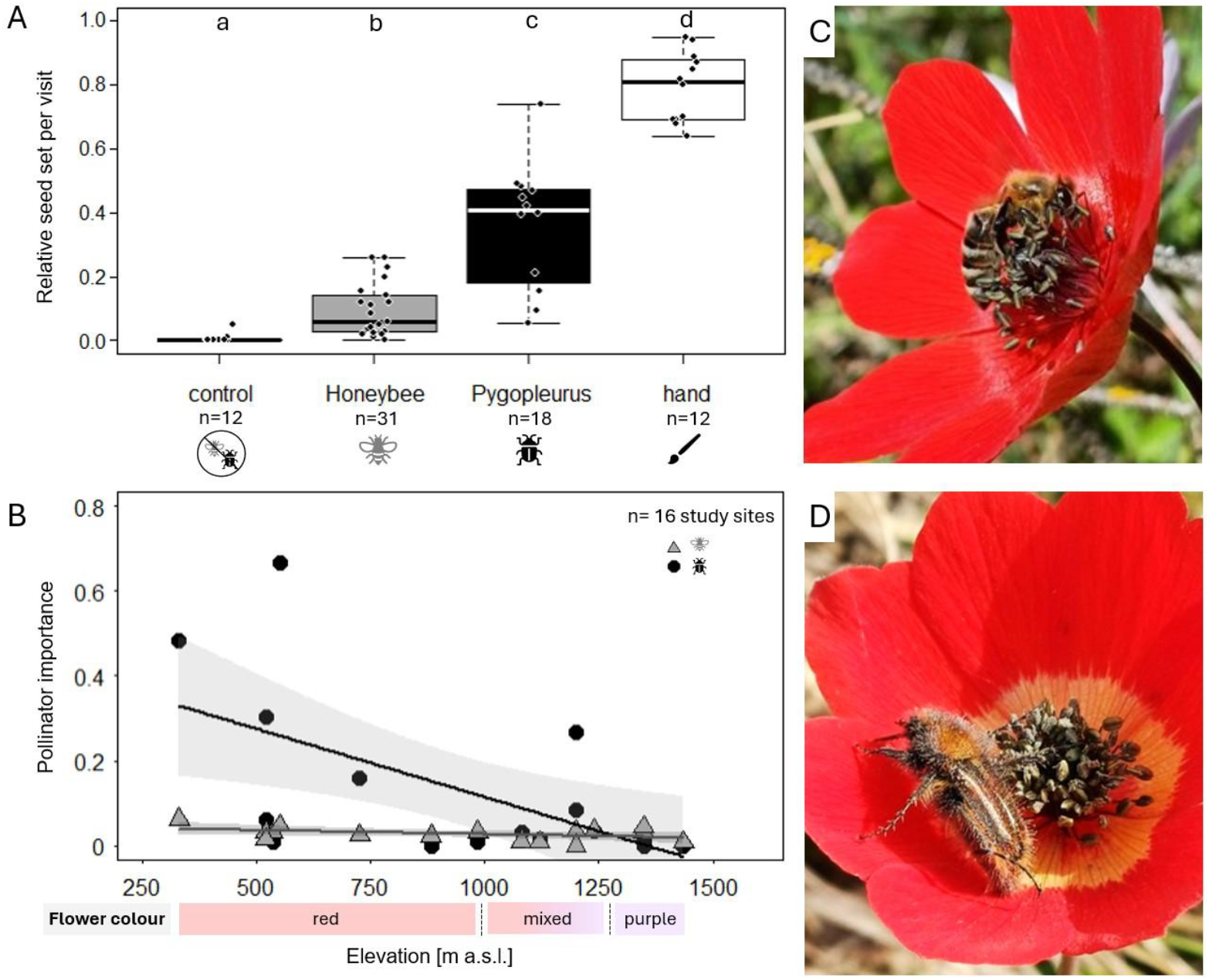
(A) Comparison between the four groups: pollinator exclusion (negative control), honeybees, *Pygopleurus* beetles, and hand pollination. The solid lines indicate the median, dots individual measurements. (B) Relationship between pollinator importance (honeybees: grey triangles; *Pygopleurus*: black dots) and elevation. The shaded area indicates the 95% CI of the regression lines (honeybees: grey; *Pygopleurus*: black). The panel on the right shows pollinators visiting red anemone flowers: (C) *Apis mellifera* and (D) *Pygopleurus chrysonotus* on *A. pavonina* flowers.

## Discussion

We compared the pollination effectiveness of bees and beetles, a largely understudied pollinator group [24], to explain geographic patterns of flower colour polymorphism in *A. pavonina*. We found differences in pollinator effectiveness of glaphyrids and honeybees, two important pollinator groups in the Mediterranean. A single beetle visit resulted in a threefold higher relative seed set as compared to a single visit by a honeybee. Consistent with their distribution along the elevational gradient [13], beetle pollinator importance declined sharply with increasing elevation, whereas that of bees remained low but constant across all elevations. On average, beetles spent considerably more time (56.3 sec.) on flowers during their visits than honeybees (2.3 sec.), which we interpret as one of the reasons for their high single-visit seed set.

Under natural (open) pollination, average seed set in the red *A. pavonina* morph is about 32% ± 6 (mean ± SE) [13], which is comparable to the observed 36.3% ± 6 seed set after beetle pollination in our experiments. We attribute the low seed set (compared to 79% in the hand pollination treatment) to pollen limitation in natural populations, likely caused by an overall low beetle abundance rather than a low pollination effectiveness of beetles. Pollen deposition generally increases with the duration of a flower visit, though the response follows a pattern of diminishing returns until it reaches an effective maximum [25, 26]. The peak of the function for some Diptera, Hymenoptera, and Coleoptera species is reached after approximately 100 seconds [26], and longer visits do not lead to additional pollen deposition. This pattern is also visible in our data (Supplementary Figure1) and indicates that *Anemone* flowers in natural population are on average visited for only about 20 seconds by beetles (corresponds to ca. 32% relative seed set). Visits longer than 100 seconds do not necessarily result in higher seed set, because Glaphyrids may also use *Anemone* flowers for mating and resting [8]. During these prolonged visits beetles likely consume pollen grains from the stigma, potentially reducing fertilization success.

The observed differences in pollination performance between beetles and honeybees may partly be caused by pollinator morphology and how pollen is transported. Pollen deposition increases with body size and hairiness of the pollinator’s body parts that touch reproductive organs [27]. Bees usually store the collected pollen on specialized body parts (e.g. corbiculae, abdominal scopa; supplementary S1A) [28] and grooming behaviour reduces the number of pollen available for outcrossing. In this context, Thomson (2003) stated that some pollinating bees resemble ‘very leaky buckets’ with respect to pollen transport and pollinator effectiveness. Whereas bees copiously groom themselves and thus remove most of the pollen from their body, we observed pollen all over beetle bodies (Figure 2D), sometimes in extremely high quantities (supplementary Figure S1B). Beetles exhibit virtually no grooming behaviour, which makes their hairy and pollen-laden bodies effective pollen vectors.

Differences in behaviour between beetles and bees inside the flower may also play a role. *Pygopleurus* beetles usually stay in the centre of the flower amidst the reproductive organs and systematically circle along the stamen to collect pollen (Supplementary Video 1A). While searching for pollen, beetles touch the reproductive organs vigorously. By contrast, honeybees are typically at the outer side of the anthers and move around the flower centre, usually with only brief, if any, physical contact with the pistils (Supplementary Video 1B). We further noticed that honeybees were more attracted to male-phase than female-phase flowers. This preference for male-phase flowers may also reduce their pollination effectiveness compared to beetles [29]. Although the importance of within-flower behaviour on the impact of pollen transport needs to be quantified, it is evident that flower-visiting behaviour is different between beetles and honeybees, and this likely contributes to their different pollinator effectiveness. The combination of morphology, foraging behaviour and the shorter *Anemone* flower visitation time renders (honey)bees lesser effective pollinators than glaphyrid beetles.

Although beetle pollination effectiveness is much higher than that of bees, the strong decline in beetle abundance with increasing elevation eventually reaches a point where the pollinator importance of beetles becomes equal to or even lower than that of bees. At this elevation, the abundance and visitation rate of other pollinators – and thus the likelihood of pollination – becomes more important. This inflection point occurs at around 1000 m a.s.l. (where the 95% CIs begin to overlap; Figure 2) and corresponds closely to the elevation at which flower colour polymorphic populations occur [12]. Thus, within the distribution range of glaphyrids, the “poppy guild” flowers are likely specialized for beetle pollination, by means of effectively excluding bees through colours that are largely inconspicuous to them [17]. We therefore conclude that flower colour variation is an adaptation shaped by geographic variation in the effectiveness and abundance of two key pollinators with different visual systems. This also neatly explains the shift from red to purple-flowered morphs at higher elevations, where only bee pollinators are present.

Honeybees are more abundant albeit less effective pollinators than solitary bee species [21, 30], some of which we have occasionally seen visiting *A. pavonina* flowers in our populations [12, 13]. Quantifying the effectiveness of solitary bees remains a challenge for future studies. However, given their high visitation rates, honeybees are one of the most common floral visitors to *A. pavonina* flowers in our natural populations [12, 13].

In summary, the higher pollinator importance (i.e. effectiveness by abundance) of beetles leads to a higher reproductive fitness at low elevations and drives the evolution of flower colours that are attractive to beetles and inconspicuous to lesser effective bee pollinators. However, this selective advantage decreases with elevation. The emergence of red-coloured, non-UV reflecting flowers at low elevations can therefore be hypothesized as an “anti-bee” strategy (bee avoidance hypothesis) [5, 31, 32]. The competitive advantage of red over purple flowers likely contributes to the maintenance of monomorphic red populations at low elevations, even though alternative pollinators (i.e. bees, which would prefer the purple morph) occur in the same area. Overall, it emphasizes the important role of pollinator-mediated selection – and differences in visual systems – as part of a continuous co-evolutionary process, with plants evolving traits to maximize reproductive fitness and match pollinator’s sensory systems.

## Funding Information

This work was supported by the Deutsche Forschungsgemeinschaft (DFG project: SP1380/2-1) to JS, the Alexander von Humboldt Foundation, Human Frontier Science Program (RGP023/2023, https://doi.org/10.52044/HFSP.RGP0232023.pc.gr.168611), and an AFOSR/EOARD grant (FA8655-23-1-7049) to CJvdK.

## Acknowledgements

We thank the Greek nature conservation authorities for the permission to our work [A.Π. ΠEN/ΔΔEΔ/32775/1175; A.Π.: YΠEN/ΔΔΔ/42915/1404; A.Π.: YΠEN/ΔΠΔ/94545/6746]; Sander de Hoop for his help with the experiments; Dimitrios Kapsalis, our contact from the local NECCA unit; and Maria and Nikolaos Mezilis and their family for their help and support in Greece.

## Notes

### Competing Interest Statement

The authors have declared no competing interest.

