## Supplementary figures and images for "Differential effectiveness of bee and beetle pollinators creates flower colour polymorphism"

### Supplementary Figure1

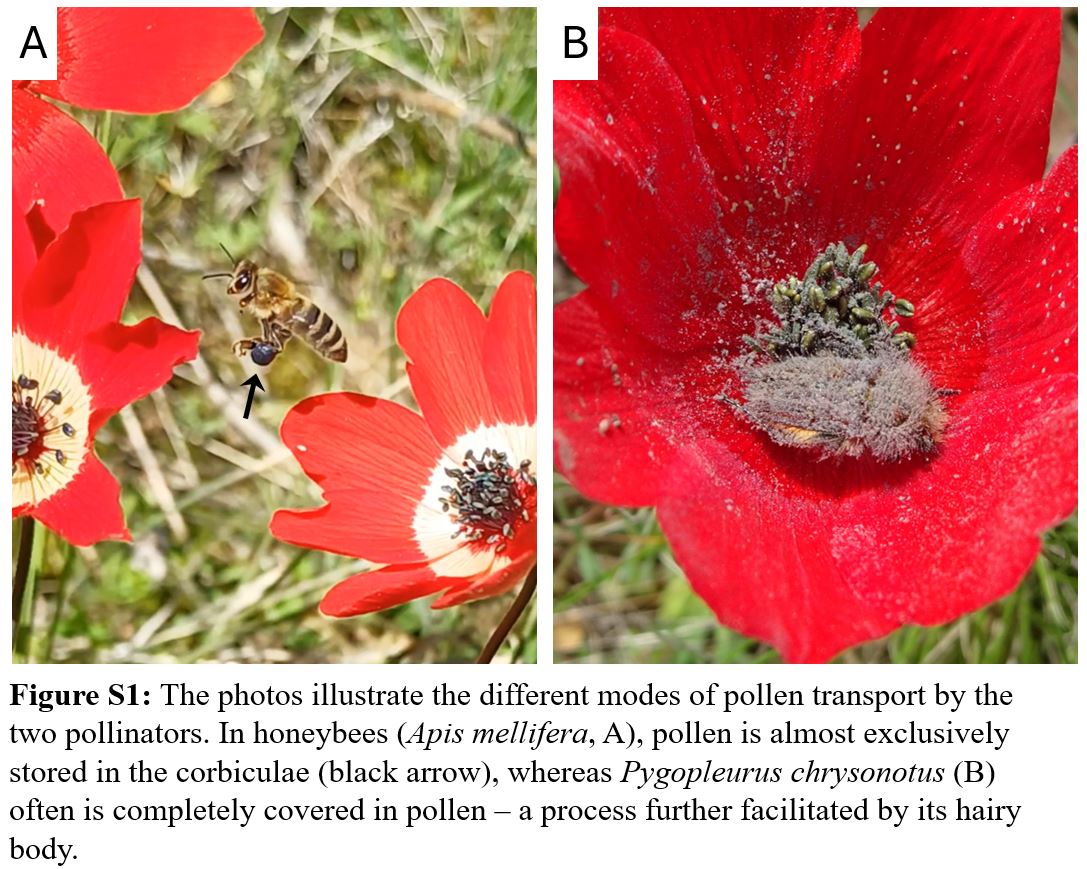

### Supplementary Figure2

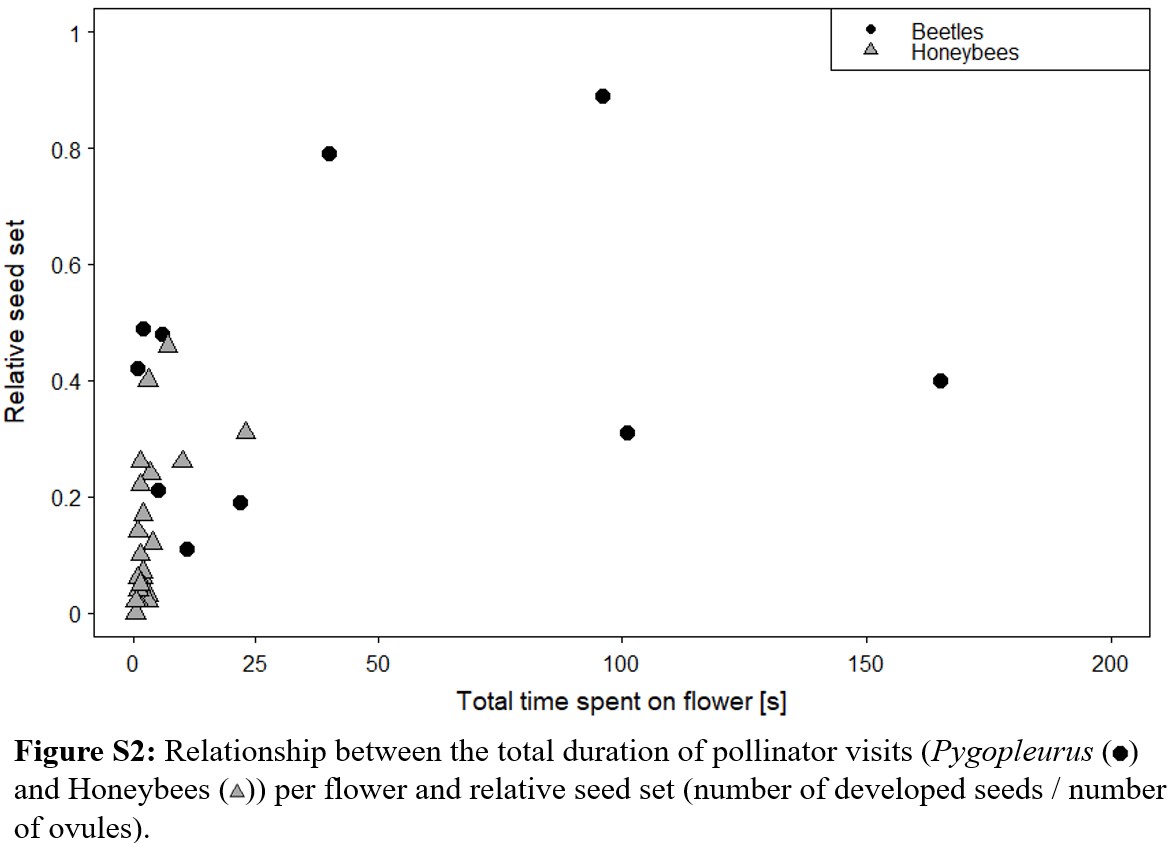
